# Modulatory Effects of the Motor Imagery and Jendrassik Maneuver upon the Forearm H-Reflex and Electroencephalographic Activity

**DOI:** 10.1101/2025.11.25.690346

**Authors:** N.M. Starodubtseva, M.V. Morozova, M.A. Lebedev

**Affiliations:** Institute for Information Transmission Problems of Russian Academy of Sciences, Moscow, Russia; Center for Neurocognitive Research (MEG Center) of MSUPE, Moscow, Russia; Lomonosov Moscow State University, Moscow, Russia

**Keywords:** H-reflex, motor imagery, Jendrassik maneuver, EEG

## Abstract

While the Jendrassik maneuver (JM) is commonly used in neurology to enhance monosynaptic reflexes such as the knee-jerk reflex, its interaction with cognitive processes like motor imagery (MI) remains unclear. In this study, we examined the effects of JM, MI, and their combination on the left flexor carpi radialis (FCR) H-reflex and cortical electroencephalographic (EEG) activity. Twenty healthy participants completed four tasks: resting, MI (imagining squeezing a tennis ball with the left hand), JM (squeezing a tennis ball with the right hand), and JM combined with MI (JM&MI). The FCR H-reflex was elicited with median nerve stimulation, and EEG changes were analyzed through event-related potentials and modulations of the mu rhythm. Surprisingly, JM performed with the right hand did not affect the contralateral H-reflex. MI of the left hand decreased the H-reflex in the right FCR. JM and MI significantly modulated cortical activity. Event-related desynchronization (ERD) of the mu-rhythm occurred during both JM and MI, and it was contralateral to the physical contraction or imagery. Bilateral ERD was observed during JM&MI. Cortical responses to nerve stimulation (P100, N100 and P300 components) varied by task, and during JM the P100 and P300 components correlated with strength of mu-rhythm ERD.

## INTRODUCTION

With the development of brain-computer interfaces (BCIs), interest has been growing to mental modulatory techniques like MI, which produces a variety of cortical modulations, including desynchronization of the mu rhythm (Llanos et al., 2013; Pfurtscheller et al., 2006). MI alone (Gentili et al., 2010; Schuster et al., 2011) and in combination with BCIs (Ma et al., 2024) has been shown to support rehabilitation in individuals with motor impairments.

The other well-known facilitation method is the Jendrassik maneuver (JM), in which a person voluntarily contracts remote muscle groups, commonly the upper limbs, such as squeezing a tennis ball or interlocking and pulling the fingers during reflex elicitation (Dowman & Wolpaw, 1988; Ertuglu et al., 2019; Tsuruike et al., 2003). Clinically, JM is routinely employed to reinforce weak or absent tendon reflexes, improving their detectability in neurological examinations. Yet, unlike MI, JM has not been accepted as a modulatory method in BCIs. Additionally, the ways in which MI and JM might interact at the cortical and spinal levels are not well understood.

In the present study, we examined the combined and separate effects of JM and MI on the forearm Hoffmann reflex (H-reflex) and associated cortical activity. The monosynaptic H-reflex has been widely used to assess spinal excitability and motor neuron pool properties (Palmieri-Smith et al., 2004). It is most commonly studied in the soleus muscle for the lower limb and the FCR for the forearm, as the reflex can be reliably elicited in these muscles at rest (Zehr, 2002). Over several decades, the H-reflex has proven to be a valuable diagnostic and research tool: alterations in reflex latency or amplitude have been associated with peripheral nerve dysfunction, changes in spinal cord excitability, and various neurological disorders (Angel, 1963; Burke, 2016).

Facilitation of the soleus H-reflex by JM has been well documented (Dowman & Wolpaw, 1988), and similar effects have been observed in the upper limb (Sugawara & Kasai, 2002). MI has been shown to significantly influence spinal excitability, as well, as demonstrated by changes in monosynaptic reflexes and responses measured using transcranial magnetic stimulation (Aoyama & Kaneko, 2011; Bonnet et al., 1997; Grosprêtre et al., 2016). To advance this research further, here we employed both JM and MI, combined with H-reflex measurements and EEG recordings, to investigate the integrative modulatory effects of these maneuvers at the cortical and spinal levels.

## MATERIALS AND METHODS

### Participants

Twenty healthy participants (12 women and 8 men; mean age and standard deviation = 23.2 ± 2.1 years; all right-handed) were involved in one experimental session each lasting 60-90 min. All participants gave written informed consent to participate in the study. The study adhered to the Human Subject Guidelines of the Declaration of Helsinki and was approved by the IITP Ethics Committee (protocol n. 2 on 16.09.2025).

### Experimental design

The experiment consisted of four tasks:

1. *Resting:* Participants remained relaxed in a sitting position without performing any overt or imagined motor activity.
2. *MI:* Participants imagined rhythmically squeezing a tennis ball with the left hand at a comfortable self-paced rhythm. They were instructed not to produce any physical movements.
3. *JM:* Participants continuously squeezed a tennis ball with their right hand.
4. *JM&MI:* Participants performed the JM task with the right hand while simultaneously performing MI of squeezing with the left hand.

The left FCR H-reflex was elicited by monopolar electrical stimulation delivered using a NeoStim-5 stimulator (Cosyma, Russia). The stimulating electrodes were positioned at the cubital fossa of the left arm, with the anode located proximally with respect to the cathode. The stimuli consisted of square-wave pulses with a duration of 1ms. They were delivered at 1 Hz. This stimulation was applied during each experimental task. Stimulus amplitude was individually adjusted for each participant to reliably elicit an H-reflex without causing discomfort.

The order of experimental tasks was randomized for each participant. Each block contained 20 randomized trials—10 task-execution trials and 10 resting trials—with each trial lasting 10 seconds. Visual stimuli specific to each condition were shown on a screen positioned 1 meter in front of the participant.

During task-execution trials, participants performed the specific task they were instructed to carry out while simultaneously counting random elements within the visual stimuli. During resting trials, participants sat relaxed, did not perform any action, and simply counted the random visual elements. Pauses were provided between experimental blocks to allow participants to rest from the stimulation before continuing.

### Data acquisition and processing

EEG data were recorded using an NVX-52 amplifier (MCS, Russia) with 22 scalp electrodes positioned according to the international 10–20 system. Ag/AgCl electrodes were lubricated with conductive gel. The ground electrode was placed at FCz. Electrode impedance was maintained below 20 kΩ throughout the recordings. Signals were sampled at 2,000 Hz.

Surface EMG activity was recorded using a bipolar electrode pair placed over the left flexor carpi radialis (FCR) muscle. The skin was prepared with alcohol-soaked cotton pads and abrasive paper to minimize contact resistance; EMG electrode impedance did not exceed 30 kΩ.

The data were processed in Python, using MNE (1.7.0) and SciPy (1.13.0). A 50-Hz notch filter was applied to remove power-line interference. The EEG signals were additionally band-pass filtered between 0.2 Hz (high-pass) and 40 Hz (low-pass) to eliminate slow drifts and high-frequency noise. Eye-movement artifacts were corrected individually for each participant using independent component analysis (ICA). The continuous EEG was then segmented into the epochs time-locked to the onset of visual stimuli, based on Lab Streaming Layer (LSL) markers.

Spectral EEG power was estimated using the Welch method, applied to each 10-s epoch. Since sensorimotor activation patterns differ between left- and right-hand movements, we compared mu-rhythm ERD in the left and right hemispheres. The activity from the C3 lead was used to express the activity of the left sensorimotor area, and the activity from the C4 lead was used for the right sensorimotor area. ERD changes relative to resting were calculated for the JM, MI and JM&MI experimental blocks using the equation:

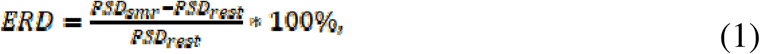

where PSD_smr_ is the power spectral density for a spectral range of interest during task execution, and PSD_rest_ is the average oscillatory power spectral density during resting within the same experimental block.

To quantify event-related potentials (ERPs), signals were re-referenced to the mastoids (A1–A2). ERP peak amplitudes were defined as average values for PO3, PO4, and Oz electrodes (for P100) and as average values for Cz and C4 electrodes (for N100 and P300).

The H-reflex amplitude was derived from the EMG trace as the peak-to-peak voltage of the response occurring shortly (10–25 ms) after the stimulus artifact.

### Statistical analysis

To account for inter-individual variability in H-reflex amplitudes, all analyses were performed on normalized data. The evoked peaks amplitudes and the power spectral density data were normalized using division by mean value across all data for each participant. First, for each participant, amplitude values for the work and rest states were sorted in ascending order to minimize the influence of random variability across stimuli. Next, each amplitude value in the work condition was divided by the corresponding value in the rest condition. This normalization step was applied separately for each participant and each condition, resulting in ratios that reflected the relative modulation of the rest state serving as an internal baseline (i.e., normalized to 1). For subsequent statistical analysis, all normalized amplitude values from all participants were combined within each condition.

Since the H-reflex data and EEG data did not meet the assumption of normality (as verified by the Shapiro–Wilk test), nonparametric tests were used. The Kruskal–Wallis test was applied to detect differences across conditions, followed by post hoc Dunn’s tests for multiple pairwise comparisons. Statistical significance was set at p < 0.05 and corrected for multiple comparisons using the Bonferroni correction where appropriate.

To examine the relationship between evoked cortical activity and sensorimotor oscillatory dynamics, a correlation analysis was performed between the amplitudes of ERP components (P100, N100, and P300) and the level of mu-rhythm ERD. Pearson’s correlation coefficients were computed between ERP peak amplitudes and the corresponding mu-ERD values separately for each hemisphere and condition.

## RESULTS

The changes in H-reflex amplitude across the conditions (JM, MI, JM & MI, and Resting) were individually variable across but small when averaged across the participants. Figure 1 presents the data for individual participants where the across-subject variability is clearly visible. Upon visual inspection, there was little to no difference in the H-reflex for the comparison of Resting with JM and JM & MI whereas one can notice an average reduction in mean the H-reflex during the MI task.

**Fig. 1.**
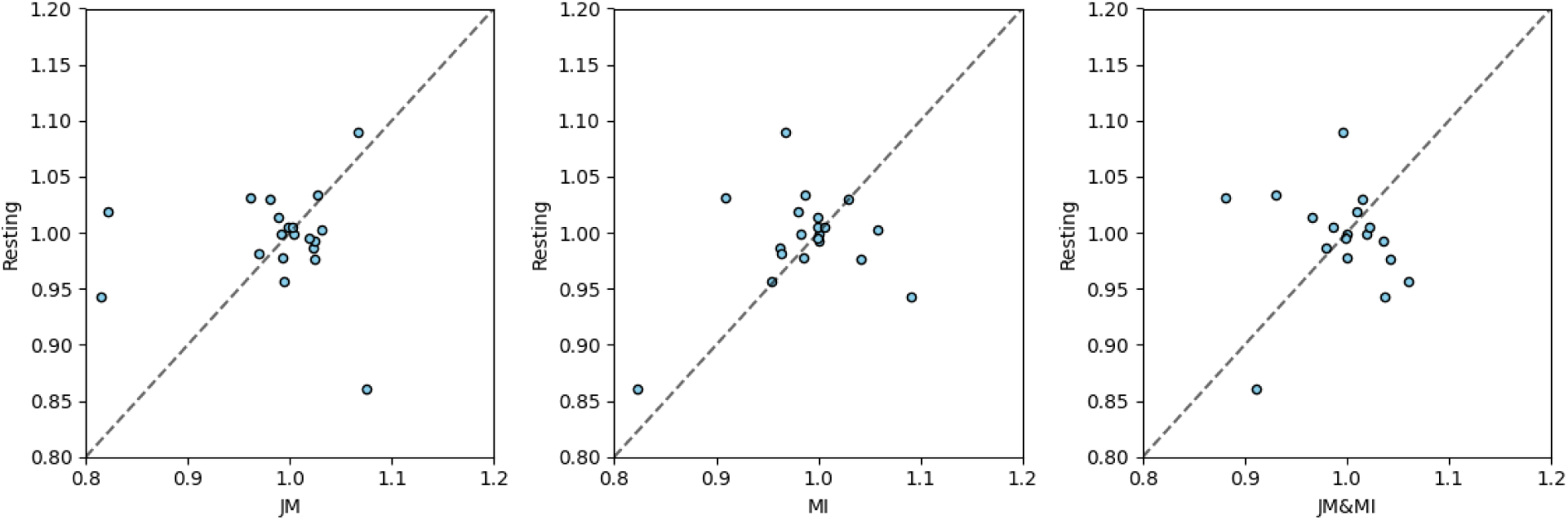
Normalized H-reflex amplitude during the JM, MI and JM&MI tasks plotted against the amplitude during Resting. Points correspond to participants.

Figure 2 shows the statistical analyses for these data. MI indeed significantly decreased the FCR H-reflex amplitude (p < 0.0001) relative to all other conditions (Fig. 2), and consistent with the visual inspection, no significant differences were found between the JM, JM & MI, and Resting conditions.

**Fig. 2.**
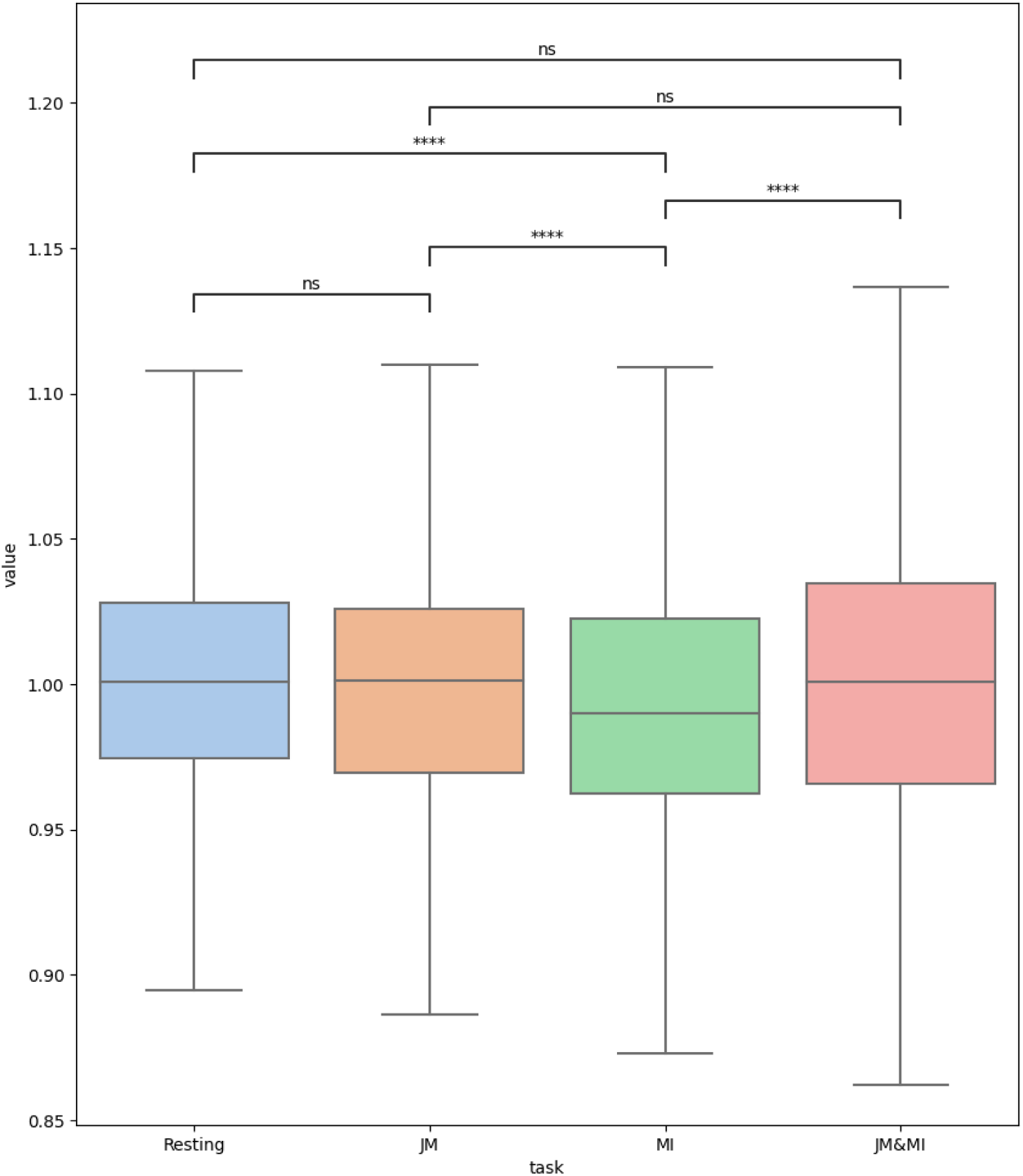
The median normalized H-reflex amplitude across all participants during Resting, JM, MI and JM&MI. Significance levels: * - p < 0.05, ** - p < 0.01, *** - p < 0.001, **** - p < 0.0001.

To evaluate EEG modulations, we first analyzed the changes in the mu-rhythm. ERD was found during JM, MI and JM & MI. The spatial distribution of mu-ERD was examined using topomaps (Fig. 3). During the JM task (where participants contracted the right hand), ERD was localized over the left sensorimotor cortex.

**Fig. 3.**
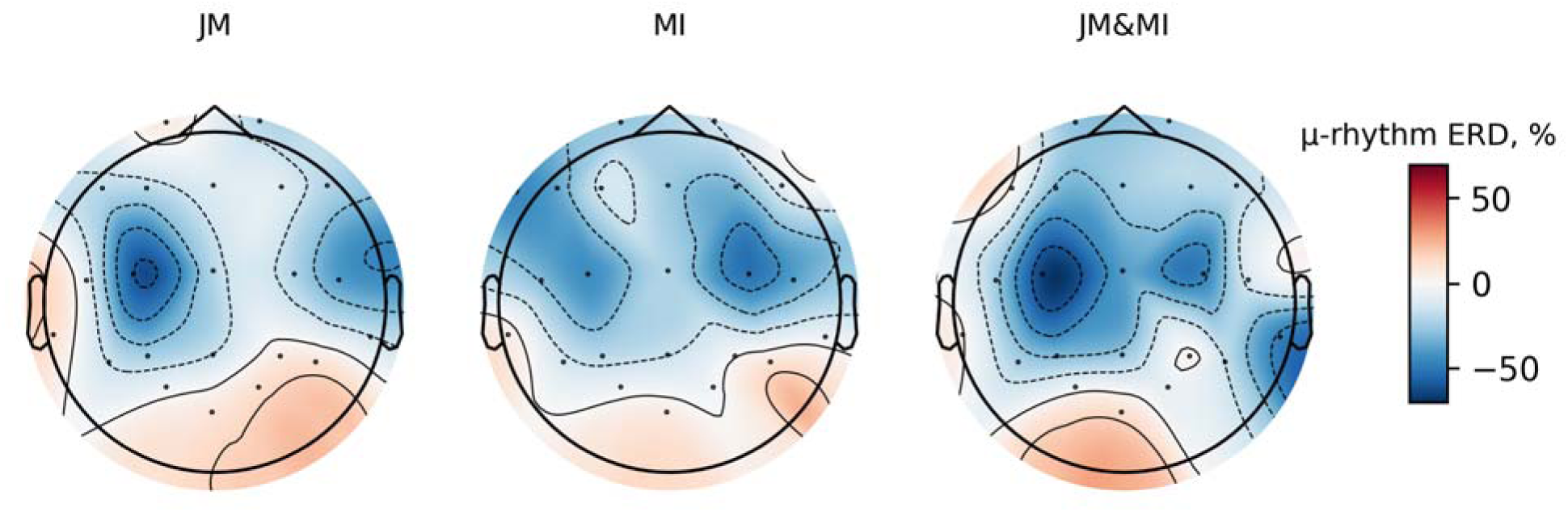
Topomaps of the mu-rhythm ERD during the JM, MI and JM&MI tasks.

During the MI task (imagining the left-hand movements), ERD occurred in both hemispheres, with a stronger effect in the right hemisphere. For the JM & MI task, bilateral activation was also evident, with a stronger ERD in the left hemisphere.

Figure 4 presents the statistical analysis of the mu-rhythm ERD in the left and right sensorimotor areas. In the left hemisphere, the strongest ERD occurred during the tasks with JM present (i.e. JM and JM & MI), and the differences in ERD were significant for the comparison of both of these tasks with MI (p < 0.05 and p < 0.001, respectively). The slight difference in ERD between the JM and JM & MI conditions was not statistically significant. In the right sensorimotor cortex, ERD was the strongest during the JM & MI condition, followed by MI, and lastly JM. All across-condition differences were statistically significant. The only significant difference between the hemispheres was the one between the mu-rhythm ERDs during JM, where ERD was clearly stronger in the left sensorimotor cortex (p < 0.0001).

**Fig. 4.**
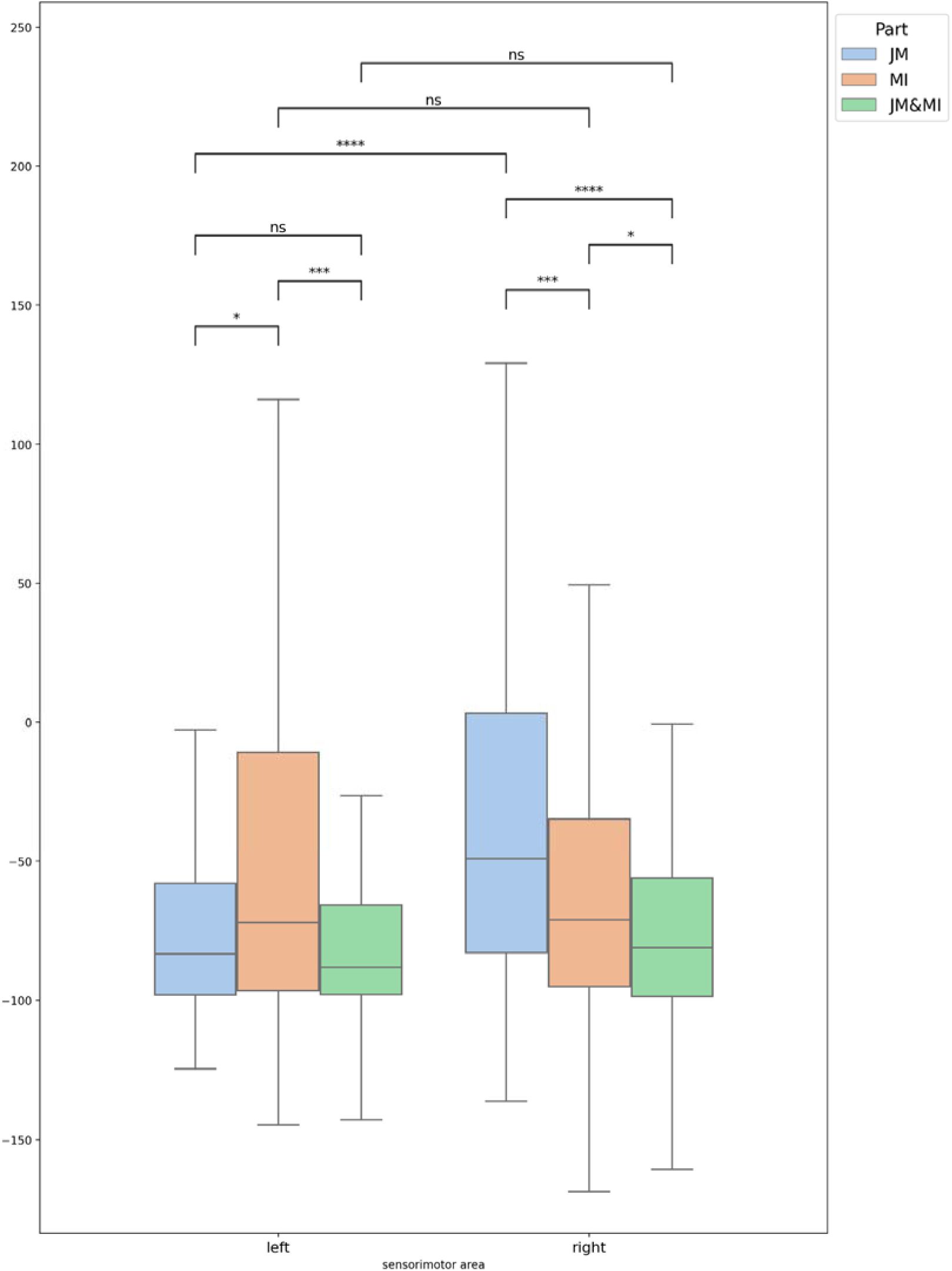
The median mu-rhythm ERD during JM, MI and JM&MI in the left and right sensorimotor areas. Significance levels: * - p < 0.05, ** - p < 0.01, *** - p < 0.001, **** - p < 0.0001.

Figure 5 shows the amplitudes of the P100, N100, and P300 components of the ERPs in response to nerve stimulation. Several differences were found across the Resting, JM, MI, and JM&MI conditions. For the P100 component, the response was stronger during JM as compared to MI (p < 0.01) and JM&MI (p < 0.01), and there were no significant differences for the other comparisons. For the N100 component, a similar trend was observed: JM facilitated the responses compared to MI (p < 0.05) and JM&MI (p < 0.05). Task-dependent effects were the strongest for the P300 component. P300 amplitude was greater during Resting as compared to MI (p < 0.0001), and JM&MI (p < 0.0001). Additionally, the P300 component was greater during JM than MI (p < 0.01) and JM&MI (p < 0.05).

**Fig. 5.**
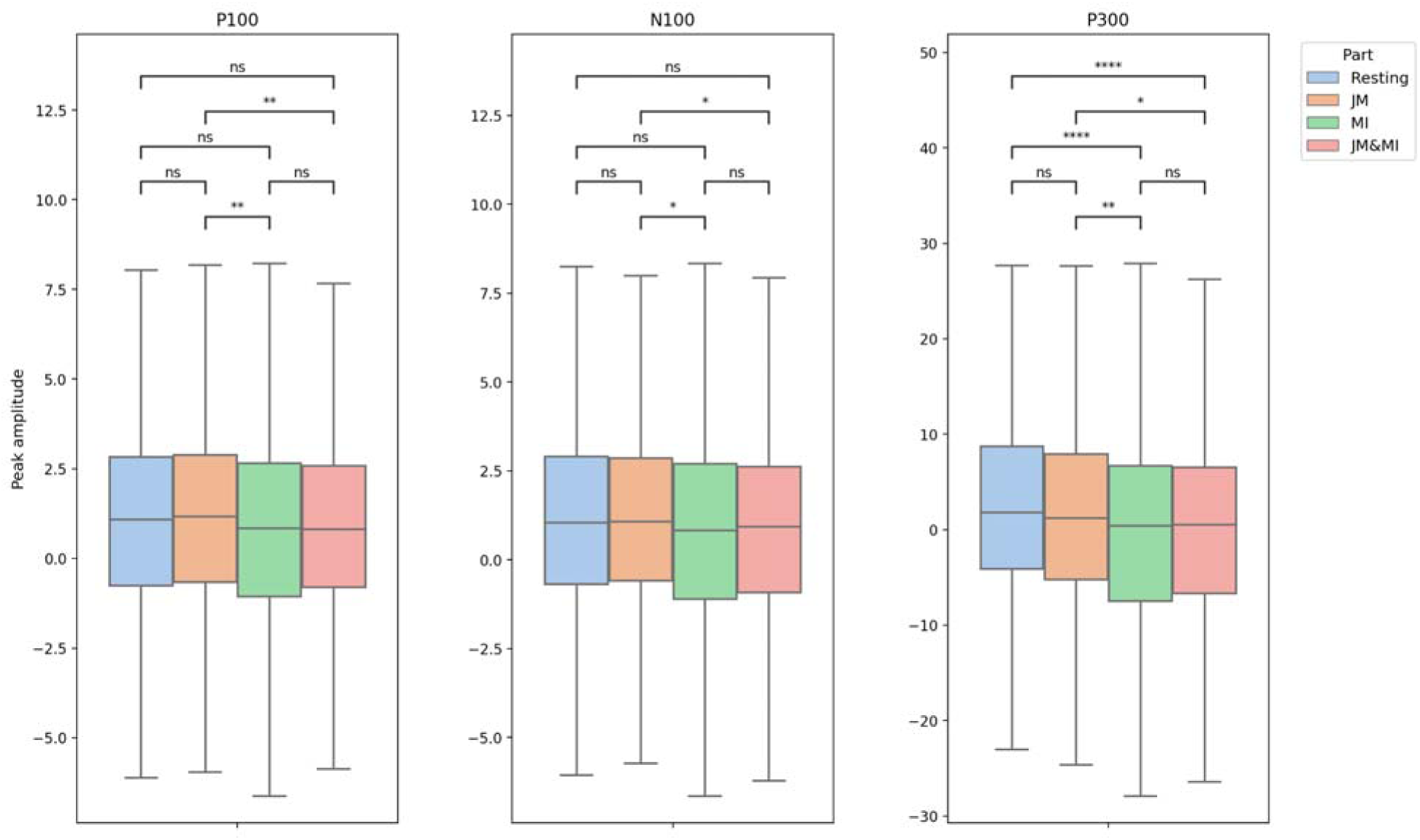
Normalized amplitudes of ERD peaks. Significance levels: * - p < 0.05, ** p < 0.01, *** - p < 0.001, **** - p < 0.0001.

The analysis of across-subject correlation analyses between the ERP components and mu-rhythm ERD showed several significant effects for different conditions. During the JM task, significant negative correlations were observed between P100 and mu-ERD for both hemispheres (left: r = −0.81, p = 1.6e-5; right: r = −0.81, p = 1.2e-5). For the same task, the P300 amplitude was negatively correlated with mu-ERD in both the left (r = −0.66, p = 1.6e-4) and right (r = −0.70, p = 6e-4) hemispheres. No significant effects were found for MI and JM & MI.

## DISCUSSION

This study examined how JM, MI, and their combination influence spinal and cortical excitability, as reflected by FCR H-reflex and EEG modulations. The findings revealed that MI decreased the H-reflex amplitude whereas JM and JM&MI did not. These results suggest that MI induces an inhibitory modulation of spinal excitability, which contrasts with the previous studies demonstrating facilitation of the H-reflex during MI of the lower limb (Hale et al., 2003) and in some cases of the forelimb (Hashemi et al., 2018).

In contrast and somewhat surprisingly, JM alone did not alter the H-reflex amplitude in the contralateral forearm, which diverges from the robust facilitation observed in the soleus (Dowman & Wolpaw, 1988). The absence of H-reflex modulation during JM&MI further indicates that JM suppresses the modulation related to MI.

Cortical activity showed clear task-related modulations. JM produced predominantly mu-rhythm ERD in the contralateral hemisphere, whereas MI induced bilateral ERD with contralateral-hemisphere dominance. The combined JM&MI condition also produced bilateral ERD, but with hemisphere dominance consistent with the stronger effect of JM (Miller et al., 2010).

EEG analyses also revealed distinct cortical modulations in response to median nerve stimulation, identified as the P100, N100, and P300 peaks. The peaks amplitude analyses revealed pronounced task-dependent differences. JM elicited larger P100 and N100 amplitudes than MI and JM&MI, reflecting enhanced early somatosensory processing during overt muscle activity (Lin et al., 2000). In contrast, the P300 component, reflecting higher-order cognitive-sensorimotor integration and stimulus evaluation (Linden, 2005), was the largest at rest and significantly reduced during MI and JM&MI. This reduction indicates that internally generated motor activity, whether imagined or combined with actual contraction, diminishes cortical responsiveness to external afferent stimuli. The P300 attenuation aligns with the inhibitory effect of MI on H-reflex amplitude, suggesting a coordinated modulation of spinal and cortical networks. Moreover, the significant negative correlations observed between P100/P300 amplitudes and mu-ERD during JM support a sensory gating mechanism, whereby desynchronization in sensorimotor rhythms during voluntary movement suppresses evoked potentials (Seki & Fetz, 2012). Interestingly, these correlations were absent in MI and JM&MI, indicating that imagery-related modulation of cortical activity operates via different top-down pathways that do not rely on afferent gating.

Overall, the results demonstrate that MI exerts inhibitory effects at the spinal level, whereas JM primarily influences cortical activity without modulating upper limbs reflexes. The absence of additive effects during JM&MI indicates that simultaneous activation of motor imagery and voluntary contraction does not simply enhance excitability but may instead produce competitive or compensatory interactions across cortical and spinal networks.

## Acknowledgements

The work was carried out as part of the state assignment of the IITP RAS, approved by the Ministry of Education and Science of Russia.

## REFERENCES

1. Angel, R. W. (1963). The H Reflex in Normal, Spastic, and Rigid Subjects. Archives of Neurology, 8(6), 591. 10.1001/archneur.1963.00460060021002

2. Aoyama, T., & Kaneko, F. (2011). The effect of motor imagery on gain modulation of the spinal reflex. Brain Research, 1372, 41–48. 10.1016/j.brainres.2010.11.023

3. Bonnet, M., Decety, J., Jeannerod, M., & Requin, J. (1997). Mental simulation of an action modulates the excitability of spinal reflex pathways in man. Cognitive Brain Research, 5(3), 221–228. 10.1016/S0926-6410(96)00072-9

4. Burke, D. (2016). Clinical uses of H reflexes of upper and lower limb muscles. Clinical Neurophysiology Practice, 1, 9–17. 10.1016/j.cnp.2016.02.003

5. Dowman, R., & Wolpaw, J. R. (1988). Jendrassik maneuver facilitates soleus H-reflex without change in average soleus motoneuron pool membrane potential. Experimental Neurology, 101(2), 288–302. 10.1016/0014-4886(88)90012-X

6. Ertuglu, L. A., Aydin, A., Kumru, H., Valls-Sole, J., Opisso, E., Cecen, S., & Türker, K. S. (2019). Jendrassik maneuver effect on spinal and brainstem reflexes. Experimental Brain Research, 237(12), 3265–3271. 10.1007/s00221-019-05668-y

7. Gentili, R., Han, C. E., Schweighofer, N., & Papaxanthis, C. (2010). Motor Learning Without Doing: Trial-by-Trial Improvement in Motor Performance During Mental Training. Journal of Neurophysiology, 104(2), 774–783. 10.1152/jn.00257.2010

8. Grosprêtre, S., Ruffino, C., & Lebon, F. (2016). Motor imagery and cortico spinal excitability: A review. European Journal of Sport Science, 16(3), 317–324. 10.1080/17461391.2015.1024756

9. Hale, B. S., Raglin, J. S., & Koceja, D. M. (2003). Effect of mental imagery of a motor task on the Hoffmann reflex. Behavioural Brain Research, 142(1–2), 81–87. 10.1016/S0166-4328(02)00397-2

10. Hashemi, S. E., Ali Ahmadi-Pajouh, M., & Shamsi, E. (2018). Does Motor Imagery Task Alter H-Reflex in FCR Muscle of The Human Hand? 2018 25th National and 3rd International Iranian Conference on Biomedical Engineering (ICBME), 1–6. 10.1109/ICBME.2018.8703590

11. Lin, Y.-Y., Simões, C., Forss, N., & Hari, R. (2000). Differential Effects of Muscle Contraction from Various Body Parts on Neuromagnetic Somatosensory Responses. NeuroImage, 11(4), 334–340. 10.1006/nimg.1999.0536

12. Linden, D. E. J. (2005). The P300: Where in the Brain Is It Produced and What Does It Tell Us? The Neuroscientist, 11(6), 563–576. 10.1177/1073858405280524

13. Llanos, C., Rodriguez, M., Rodriguez-Sabate, C., Morales, I., & Sabate, M. (2013). Mu-rhythm changes during the planning of motor and motor imagery actions. Neuropsychologia, 51(6), 1019–1026. 10.1016/j.neuropsychologia.2013.02.008

14. Ma, Z.-Z., Wu, J.-J., Cao, Z., Hua, X.-Y., Zheng, M.-X., Xing, X.-X., Ma, J., & Xu, J.-G. (2024). Motor imagery-based brain–computer interface rehabilitation programs enhance upper extremity performance and cortical activation in stroke patients. Journal of NeuroEngineering and Rehabilitation, 21(1), 91. 10.1186/s12984-024-01387-w

15. Miller, K. J., Schalk, G., Fetz, E. E., den Nijs, M., Ojemann, J. G., & Rao, R. P. N. (2010). Cortical activity during motor execution, motor imagery, and imagery-based online feedback. Proceedings of the National Academy of Sciences, 107(9), 4430–4435. 10.1073/pnas.0913697107

16. Palmieri-Smith, R., Ingersoll, C., & Hoffman, M. (2004). The Hoffmann Reflex: Methodologic Considerations and Applications for Use in Sports Medicine and Athletic Training Research. Journal of Athletic Training, 39, 268–277.

17. Pfurtscheller, G., Brunner, C., Schlögl, A., & Lopes da Silva, F. H. (2006). Mu rhythm (de)synchronization and EEG single-trial classification of different motor imagery tasks. NeuroImage, 31(1), 153–159. 10.1016/j.neuroimage.2005.12.003

18. Schuster, C., Hilfiker, R., Amft, O., Scheidhauer, A., Andrews, B., Butler, J., Kischka, U., & Ettlin, T. (2011). Best practice for motor imagery: a systematic literature review on motor imagery training elements in five different disciplines. BMC Medicine, 9(1), 75. 10.1186/1741-7015-9-75

19. Seki, K., & Fetz, E. E. (2012). Gating of Sensory Input at Spinal and Cortical Levels during Preparation and Execution of Voluntary Movement. The Journal of Neuroscience, 32(3), 890–902. 10.1523/JNEUROSCI.4958-11.2012

20. Sugawara, K., & Kasai, T. (2002). Facilitation of motor evoked potentials and H-reflexes of flexor carpi radialis muscle induced by voluntary teeth clenching. Human Movement Science, 21(2), 203–212. 10.1016/S0167-9457(02)00099-4

21. Tsuruike, M., Koceja, D. M., Yabe, K., & Shima, N. (2003). Age comparison of H-reflex modulation with the Jendrássik maneuver and postural complexity. Clinical Neurophysiology, 114(5), 945–953. 10.1016/S1388-2457(03)00039-7

22. Zehr, E. P. (2002). Considerations for use of the Hoffmann reflex in exercise studies. European Journal of Applied Physiology, 86(6), 455–468. 10.1007/s00421-002-0577-5

